# Maternal Continuous Oral Oxycodone Self-Administration Alters Pup Affective/Social Communication but not Spatial Learning or Sensory-Motor Function

**DOI:** 10.1101/2020.04.04.022533

**Authors:** Giulia Zanni, Patrese A. Robinson-Drummer, Ashlee A. Dougher, Hannah M. Deutsch, Matthew J. DeSalle, David Teplitsky, Aishwarya Vemulapalli, Regina M. Sullivan, Amelia J. Eisch, Gordon A. Barr

**Affiliations:** Department of Anesthesiology and Critical Care Medicine, Children’s Hospital of Philadelphia and University of Pennsylvania, Philadelphia, PA, USA; Department of Child and Adolescent Psychiatry, New York University Langone Medical Center, New York, NY, United States; Emotional Brain Institute, Nathan Kline Institute for Psychiatric Research, Orangeburg, NY, United States; Department of Neuroscience, Perelman School of Medicine - University of Pennsylvania, Philadelphia, PA; Department of Psychology, University of Pennsylvania, Philadelphia, PA

**Author notes:** Correspondence can be addressed to either or both. AJE, GAB. Co-first authors, equal contribution (Giulia Zanni, Patrese A. Robinson-Drummer). GZ: Department of Developmental Neuroscience, New York State Psychiatric Institute, New York, NY, PR-D: Department of Psychology, Haverford College, Haverford, PA. Co-senior authors, equal contribution (Amelia J. Eisch, Gordon A. Barr).

## Abstract

The broad use and misuse of prescription opioids during pregnancy has resulted in a surge of infants diagnosed with Neonatal Opioid Withdrawal Syndrome (NOWS). Short-term irritability and neurological complications are hallmarks of NOWS, but the long-term consequences are unknown. Our newly-developed preclinical model of oxycodone self-administration enables adult female rats to readily drink oxycodone (0.06-0.12 mg/ml, ∼10/mg/kg/day) continuously before and during pregnancy and after delivery, to achieve similar liquid intake in oxycodone moms relative to water-only controls. Oxycodone levels were detected in the serum of mothers and pups. Growth parameters in dams and pups, and litter mass and size were similar to controls. Maternal behavior at postnatal day 1 (PN1) was unchanged by perinatal oxycodone consumption. Regarding the plantar thermal response, there were no differences in paw retraction latency between oxycodone and control pups at PN2 or PN14. Oxycodone and control pups had similar motor coordination, cliff avoidance, righting time, pivoting, and olfactory spatial learning from PN3 through PN13. Separation-induced ultrasonic vocalizations at PN8 revealed higher call frequency in oxycodone pups relative to controls. Finally, during naltrexone precipitated withdrawal at PN9, oxycodone males vocalized more than control pups, consistent with a previously-published withdrawal phenotype. Thus, our rat model of continuous oral oxycodone self-administration in pregnancy shows exacerbated affect/social communication in pups in a sex-dependent manner but spared cognition and locomotion. Our preclinical, high face validity NOWS model reproduces key aspects of human opioid use during pregnancy, enabling longitudinal analysis of how maternal oxycodone changes emotional behavior in the offspring.

**HIGHLIGHTS:**

- Female rats self-administered oxycodone at clinically relevant doses before and during pregnancy and for the first two weeks after parturition.
- Both dams and pups, for the14 day postnatal experimental period, had detectable levels of oxycodone in their blood
- Dams drinking oxycodone only or water only did not differ in weight gain, water intake, or the number of pups born and their pups did not differ in weight throughout.
- Sensory and motor function in the pups was not altered, nor was hippocampal dependent spatial learning.
- Oxycodone exposed pups were physically dependent and displayed increased withdrawal behaviors with or without the opioid antagonist naltrexone.
- Pups expressed more negative affect, expressed by increased ultrasonic vocalizations, following naltrexone precipitated withdrawal or when separated from their mother.

## Introduction

The growing opioid epidemic has led to increased opioid use during pregnancy, resulting in a nearly five-fold increase in newborns exposed to opioids^1^. These newborns experience neonatal opioid withdrawal syndrome [NOWS^2,3^], showing irritability, high-pitched crying, gastrointestinal and autonomic dysregulation and longer hospitalizations^3,4^.

Preclinical opioid exposure models enable controlled, single-substance approaches to identifying opioid-related neurobehavioral alterations and causative mechanisms, overcoming the confounding factors inherent in human studies^5^. Some studies employ repeated, once or twice daily parenteral injections of opioids to the mother^6-10^. Other studies use continuous administration via minipumps or pellets ensuring a steadier release of the opioid. But because the dam’s weight increases throughout gestation, the dose decreases proportionately^11-15^. Both models inherently include fetal *in utero* exposure and withdrawal from opioids due to spontaneous withdrawal or the termination of treatment at parturition^16^. Additionally, the mother may be stressed if treatment begins during gestation, causing increased or irregular corticosterone release, making it challenging to discern direct drug effects from those of hypothalamic-pituitary-adrenal activation^17-19^. Injections of morphine given to pups postnatally avoid the confounds related to maternal physiological changes, but not withdrawal, and only model drug exposure during the equivalent of the human third trimester, reducing their translational relevance^20,21^.

These models contribute substantially to our knowledge of NOWS. Here we expand upon this research using oxycodone, a drug at the center of the rising opioid epidemic^1^. Rodent models of oxycodone use employ once or twice daily oral gavage or short-term, intravenous self- or experimenter-administration^5,22-25^. This early exposure is associated with hyperactivity in adult animals^25^, altered stress responsivity^24^ and disrupted spatial learning^22^, recapitulating some of the long-term effects observed in humans^26^. Recently, we developed an oral oxycodone self-administration protocol in which rats have access to and freely drink oxycodone, escalate their intake over months, are highly-motivated to continue intake, and become physically dependent, emulating the human profile of opioid use disorder^27^. Nonetheless, oral oxycodone self-administration has not been assessed in pregnant dams^28,29^. Here we apply this protocol before, during, and after pregnancy, allowing dams to titrate their intake, avoiding spontaneous withdrawal^27^. Furthermore, as dams ingest oxycodone prior to breeding and through lactation, developing pups are exposed to the drug during all phases of fetal and infant development, making it a valuable tool for modeling NOWS following neonatal opioid exposure.

To elucidate the effects of ante- and early post-natal exposure to oxycodone, we screened a constellation of neurodevelopmental, affective, and cognitive alterations in infants to characterize NOWS. Our results show that following perinatal oxycodone exposure, drug-exposed pups have typical sensory-motor development and spatial learning relative to water-control pups; however, drug exposure altered affective states following maternal separation and precipitated drug withdrawal.

## Methods

### Ethical permission guidelines

All procedures were conducted in accordance with the NIH Guide for the Care and Use of Laboratory Animals (NIH Publications No. 80-23) and the guidelines of the Society for Neuroscience and the International Society of Developmental Psychobiology. All methods were approved in advance by the Animal Care and Use Committee of the Children’s Hospital of Philadelphia. Our protocol included defined humane endpoints for euthanizing animals. No animal reached these humane endpoints prior to experimental endpoints.

### Subjects

Subjects were adult male and female Hooded Long-Evans rats purchased from Envigo and their offspring. Three separate cohorts were tested over the course of two years. Rats for cohorts 1-2 arrived at 70-75 days of age; rats for cohort 3 arrived at 50 days old. On arrival, male rats were housed singly and female rats housed in pairs for 10-14 days with *ad libitum* food and water (cage size: 51×40cm) in a temperature-controlled vivarium (20-23°C and humidity 30-70%), 12:12 photocycle, lights on at 0600h. Females were then individually-housed until breeding. Rats were weighed and handled three times/week for 2-3 minutes to familiarize them to different experimenters. Cages were cleaned once a week with an occasional spot-cleaning as-needed.

At the start of the experiment, prior to breeding, females were acclimated to water-only by placing either a Hydropac® (cohorts 1 and 2) or Zyfone bottle (cohort 3) in each of two water hopper ports within the cage. Following acclimation, females were randomly assigned to the oxycodone only (0.06-0.12 mg/mL; Experimental) or water (Control) group. Liquid intake was measured before breeding, throughout breeding, and during gestation (Figure 1). This concentration of oxycodone elicited stable self-administered doses of ∼10 mg/kg/day (Figure 1) prior and subsequent to breeding.

**Figure 1:**
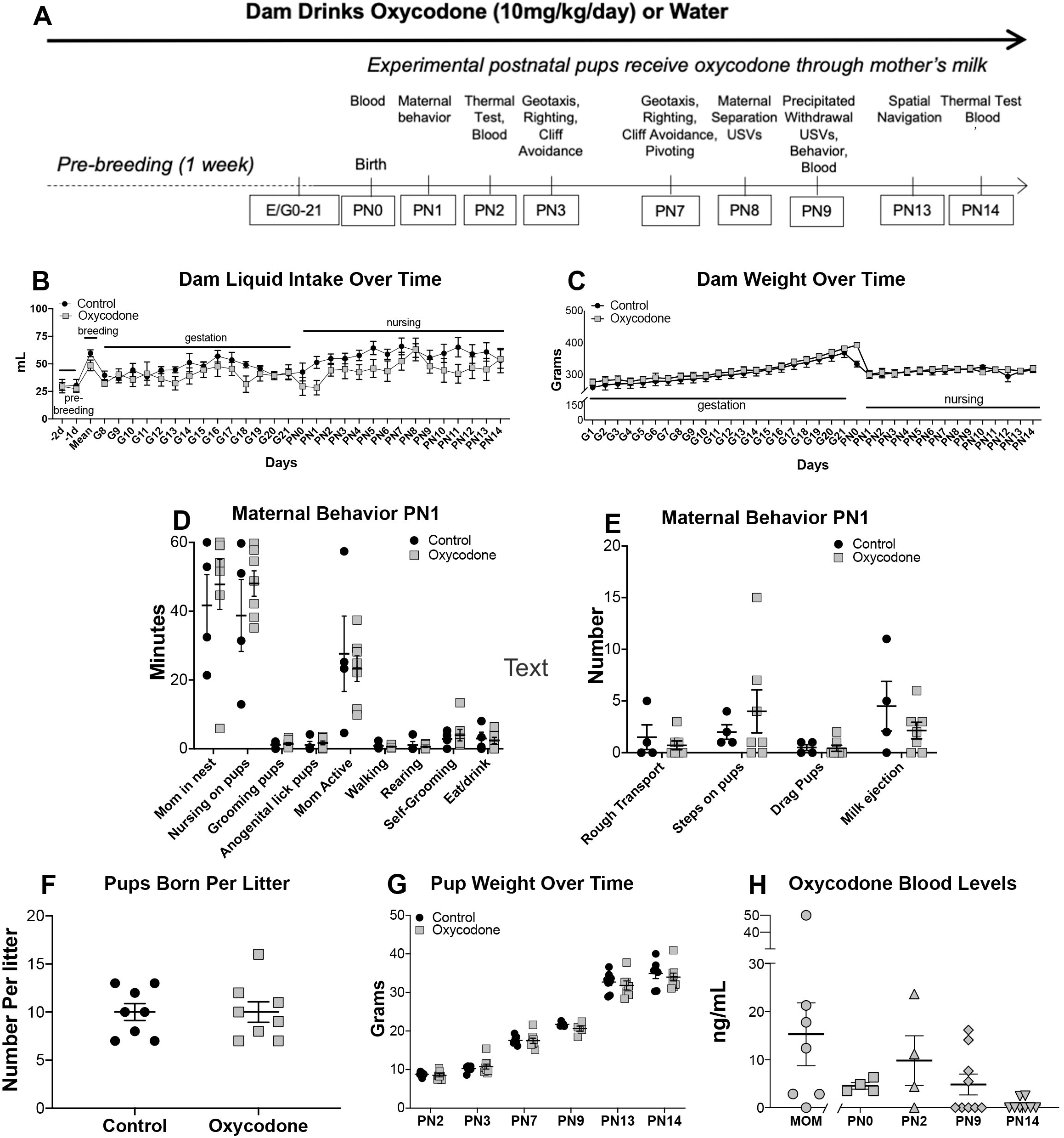
Maternal intake, growth, and behaviors. **A.** Timeline depicting course of pup pre/postnatal oxycodone or control exposure and experimental procedures. **E**, embryonic day; **G**, gestational day; **PN**, postnatal day; **USV**, ultrasonic vocalizations. **B.** Maternal liquid intake for oxycodone and control dams increased overtime in both groups (*p* < 0.001) with no difference between groups (*p =*0.707). **C.** Dam weight increased similarly during gestation (G1-G21, *p* < 0.001) and lactation (PN0-PN14, *p* < 0.001) with no difference between groups (p *=* 0.575). **D.** Mean ± SEM maternal and self-care behavior total duration observed on PN1. No difference was observed between groups (all *p*’s > 0.050). **E.** Mean ± SEM rough behavior and milk ejection total frequency observed on PN1. No difference was observed between groups (all *p*’s > 0.050). **F.** Mean ± SEM number of pups born per litter to oxycodone and control litters. There was no difference in litter sizes, *p* = 0.999). **G.** Mean ± SEM pup weight from PN2-PN14 in oxycodone-exposed and control litters; weight increased significantly (*p* < 0.001) but no difference in weight gain was observed between groups [p *=* 0.605). **H.** Mean blood levels of oxycodone in dams and pups at PN0, PN2, PN9, and PN14.

One week after oxycodone/water administration started, a male rat was introduced to the cage of the female. In cohort 1, one male and one female were placed together for 4 to 18 days, in cohorts for 7 days, and for cohort 3 for 4 days. After breeding, dam weights were recorded daily. After birth, pup weights were recorded prior to each behavioral test. Corn cob bedding was supplemented with non-toxic shredded paper for nest building and a plastic enrichment tunnel.

Cages were checked for pups twice daily, ∼0900h and 1700h. The day of birth was designated postnatal day (PN) 0 with no cage changes until 2 days after delivery. On PN2, litters were culled to 8-10 pups with equal males and females when possible.

### Maternal behavior

Maternal behavior of the dam with the pups was assessed using previously-validated behavioral observation protocols^30,31^ (Table 1). On PN1 and PN2, video cameras were directed toward the broad side of the cage and maternal activity was recorded for 60 minutes beginning at 10:00h. Blind scoring of maternal behavior at PN1 was performed by 2 independent observers offline. Scoring of data from PN2 was by a single blind observer, also offline. Only the PN1 data are reported here.

**Table 1:** List and description of behaviors recorded and analyzed during maternal behavior observations

|  | Behavior | Description |
| --- | --- | --- |
| *Maternal Care* | Nursing on pups | Mom is hunched over actively, or laying on her side passively, with pups attached to nipples |
|  | Grooming pups | Mom cleaning body of pup with her tongue |
|  | Anogenital lick pups | Mom is using tongue to stimulate anogenital region of pup |
| *Nesting Behavior* | Pups all in nest | All pups gathered in nest huddle |
|  | Mom in nest | Mom's entire body is inside nest area |
|  | Nest Building | Mom is actively manipulating bedding material in or around the nest (i.e. shredded paper) |
| *Dam Activity* | Mom Active | Any instance of activity where mom is moving inside or outside of the nest |
|  | Walking | Mom is walking around the cage |
|  | Eat/drink | Mom is eating food pellets or drinking from water container |
|  | Rearing | Mom is sitting on back paws with or without her paws resting on the cage wall |
|  | Self-Grooming | Mom is using her tongue to clean her body |
| *Rough handling* | Rough transport | Picking up and moving a pup by any area that is not the nap of the neck |
|  | Stepping on pups | Mom tramples pups under foot |
|  | Dragging pups | While pups are still nipple-attached, mom walks away from nest dragging pup underneath |
| *Milk Ejections* |  | While nipple-attached, milk is released causing pups to reflexively elongate their bodies and become momentarily active |

**Note:** Behaviors were compared between moms exposed to oxycodone before, during, and after gestation and control moms exposed only to water.

### Dam and pup oxycodone blood levels

Blood was collected from the dams and male and female pups in different litters to assay oxycodone levels. When needed, blood was pooled from same-sex pups within litters. Trunk blood was taken via rapid decapitation for pups at PN0, PN2, PN9, and PN14, and 14 days after delivery for dams. For PN0, PN2, and PN14, blood was captured between 1000h–1700h, the day part of the light cycle. For PN9, blood was taken immediately after the naltrexone-precipitated withdrawal test. The blood was centrifuged at 4K RPM, 4°C, for 10 minutes, serum separated, stored at -20°C, until sent to the Bioanalytical Core Center for Clinical Pharmacology at the Children’s Hospital of Philadelphia Research Institute for analysis. Assay development has been described previously^27^.

### Plantar thermal test

PN2 and PN14 pups were tested for the latency to withdraw their hindpaws to a thermal stimulus (Plantar withdrawal test; “Hargreave’s” apparatus; IITC Inc. Life Science, Model 390G, Heated Base) as previously described^27,32^. Pups were placed singly within an inverted plexiglass cage on an elevated glass surface maintained at 30°C. Following 10 minutes of acclimation, a radiant heat source beneath the glass converged on the plantar surface of the hindpaw and the latency to withdraw measured. Cut-off time was set at 20 seconds to avoid tissue injury. The intensity of the light was set to provide a withdrawal response of about 6-8 seconds at PN2 and 10-12 seconds at PN14, allowing either increased or decreased response latencies. Each hindpaw was tested three times and withdrawal latencies were averaged across trials and within litters to produce a single litter score for each sex.

### Neurodevelopmental Tests

Pups were tested for the behaviors in the order listed below between 1000h and 1200h at PN3 and PN7, except for pivoting tested which was performed only at PN7. Tests were adapted from previous work^33,34^. The entire litter and dam were transported to the testing room. Pups removed from the dam, individually weighed and held as a group in a cage with nesting material on a heating pad (30°C). Each test was done three times in rapid succession and scores averaged. Scores from individual pups were averaged within litters for each sex. Each pup tested took about 6-8 minutes and the litter was away from the dam for 45-60 minutes.

#### Negative Geotaxis

Pups were placed head down on a mesh surface tilted at a 20° angle. The latency to make a full 180° turn was measured. Trials were capped at 60 seconds.

#### Surface Righting

Pups were placed on their backs on a warmed heavy black plexiglass sheet (30-32°C) and the latency to turn over with all four paws touching the surface was measured. Cutoff latencies were set at 30 seconds.

#### Cliff Avoidance

Pups snout and forepaws were extended over the edge of an elevated surface and latency to move their head and paws away from the edge within 30 second recorded.

#### Pivoting

Pups were placed on the warmed heavy black Plexiglas sheet and the number of 90° turns executed within 60 seconds recorded.

### Maternal separation ultrasonic vocalizations

Separation-induced ultrasonic vocalizations (USVs) were measured individually at PN8^35^. Pups were weighed and placed in a square container (19×33cm) filled with clean cage bedding inside a 32°C air-jacketed incubator (Jeio Tech, Model IB-05G). Vocalizations were captured using a Dodotronics 200kHz USB microphone (Castel Gandolfo, Italy) and analyzed using Raven Pro (version 64 1.4) software. Using standard measures provided by Raven Pro, vocalizations were assessed for total call count, USV peak frequency (kHz), and USV peak power (dB). After five minutes of separation, pups were returned to the mom for two minutes. A second five-minute separation and USV recording were performed.

### Precipitated withdrawal

PN9 pups were weighed and injected intraperitoneally with naltrexone HCl (Sigma; 1mg/kg, i.p.; 0.1mg/mL) dissolved in saline, or saline alone. Two males and two females from each Control or Experimental litter were individually weighed, injected, and immediately placed in the incubator. USV’s were recorded for five minutes (see Maternal separation USVs^35,36^). Video recordings of pup behavior were analyzed blindly for specific withdrawal behaviors (Table 2). Following USV and video recording, all injected pups were sacrificed by rapid decapitation and trunk blood collected in an Eppendorf tube and promptly put on ice.

**Table 2:**
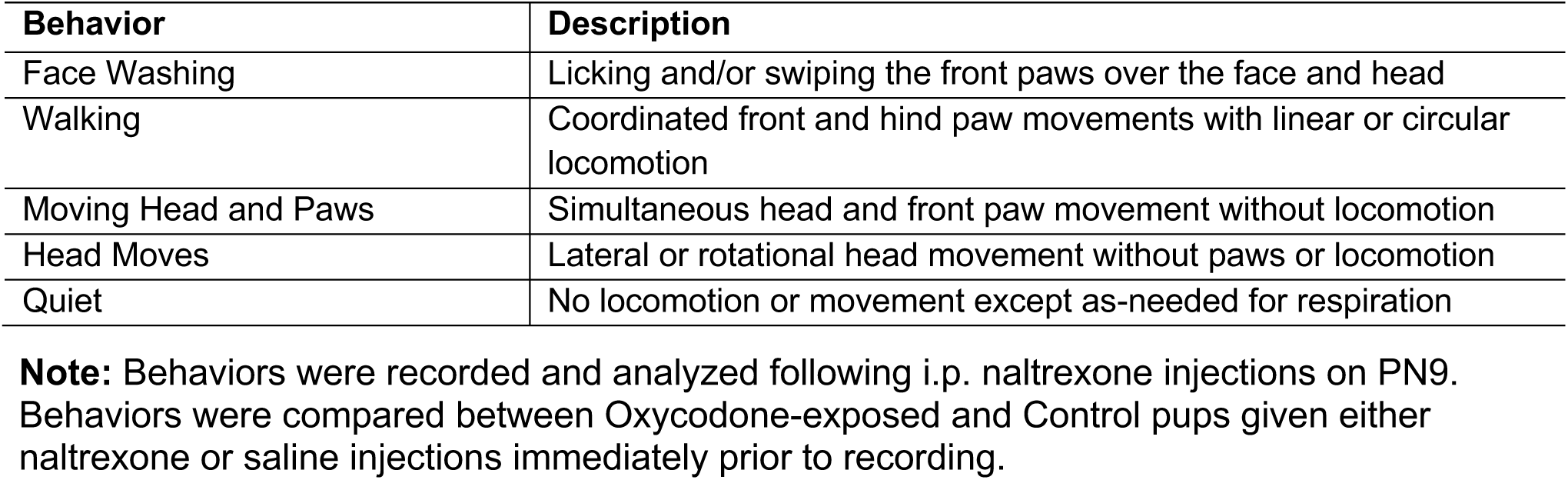
List and description of pup withdrawal behaviors.

### Spatial navigation

PN13 pups were tested for hippocampus-dependent spatial learning^37,38^ using odor cues. Pups were acclimated for two minutes to plastic warmed opaque goal box (8×12cm) that held pups and soiled bedding within an arena (45×32cm) with 1cm of clean cage bedding. The goal box was placed lateral and away from the center of the arena during training (Figure 3B). There were four training trials for pups to learn to navigate to the goal box followed by a single test trial in which the goal box was removed and replaced with an empty goal box. The time to locate the new box was measured. For training, one pup was removed from the huddle in the goal box, placed facing the wall in one of the two arena corners most distant from the goal box, and allowed to freely navigate the arena. Acquisition trials were terminated when a pup contacted the goal box or after 150 seconds. The pup was then returned to the huddle. This procedure was performed four times, and after each trial the bedding was thoroughly mixed, smoothed, and the pup repositioned at the alternate starting position. After the fourth trial, littermates were removed, the goal box replaced, and the test trial began. Pups were placed in the corner opposite the last acquisition trial and the latency to enter the former location of the littermate box was recorded (cutoff time was 180 seconds). To serve as navigation cues, a cotton applicator tip fixed 8 cm above the floor in each corner of the arena was infused with 100uL of either anise oil, coconut oil, or orange or peppermint extract (all from McCormick). There were two configurations: the “Fixed” procedure in which all cues remained constant between trials and the “Alternating” procedure in which the four odors were randomly exchanged among the corners after each trial avoiding the same configuration twice, eliminating the use of these cues to navigate to the goal. Within a litter, half of the pups received the Fixed training and half received the Alternating training. Cohort 3 was tested in a different room that had inadequate ventilation such that the odors permeated the room and precluded valid test results. Therefore the spatial navigation data presented are only from cohorts 1 and 2.

**Figure 2:**
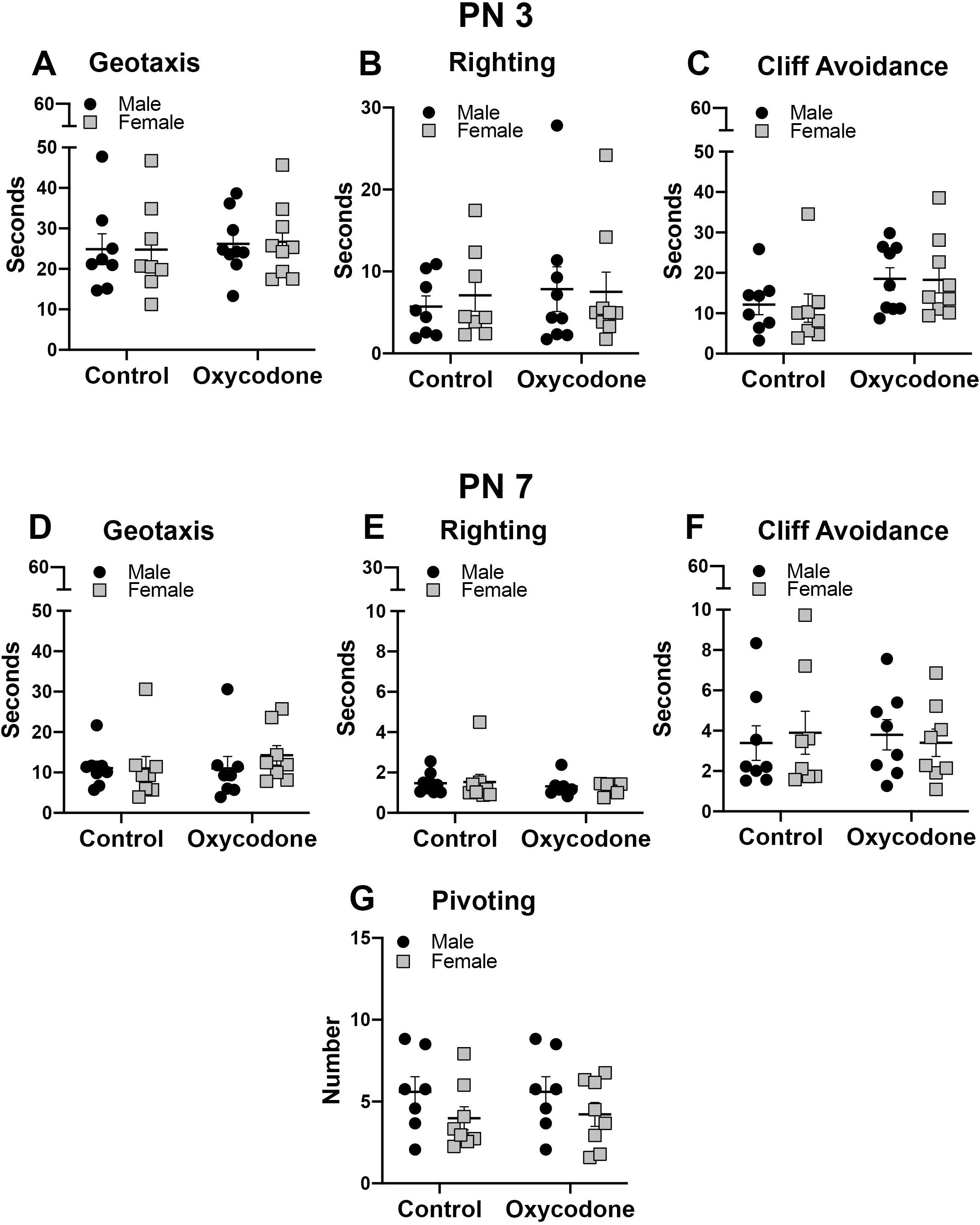
Neurodevelopmental tests performed on (top) PN3 or (bottom) PN7. There was no difference observed between groups in any test (all *p*’s > 0.050). Geotaxis: Mean ± SEM seconds to reorient head and body on a 20° slope in 60 seconds. Righting: Mean ± SEM seconds to rotate from supine to prone position within 30 seconds. Cliff Avoidance: Mean ± SEM seconds to move backwards from an elevated edge in 30 seconds. Pivoting: Mean ± SEM total frequency of 90° turns in 60 seconds.

**Figure 3:**
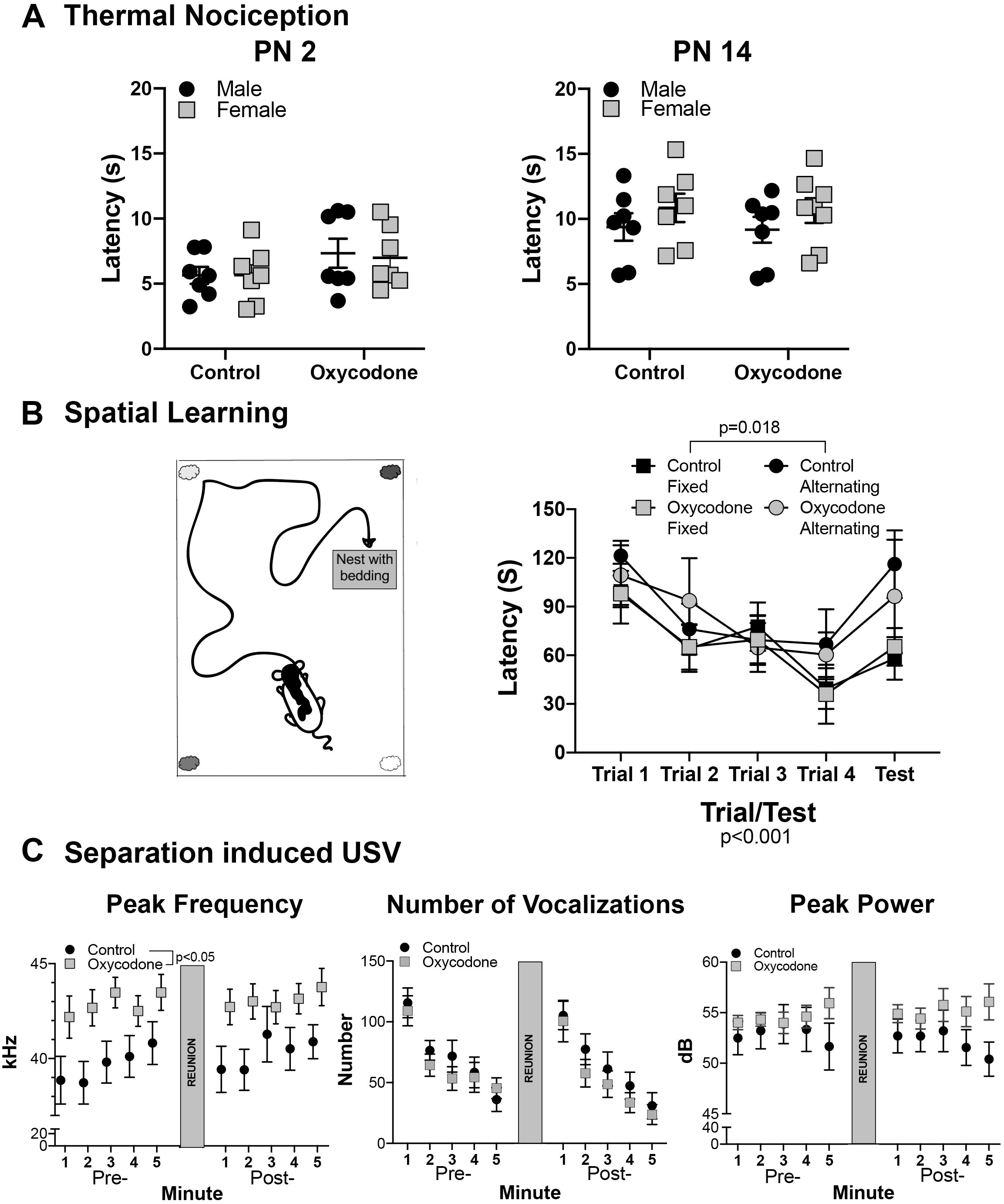
Thermal nociception tests, spatial learning, and separation induced USVs. **A.** Mean± SEM latency (seconds, s) to withdraw hindpaw from noxious thermal stimulus on PN2 (left graph) or PN14 (right graph). There was no difference in withdraw latencies observed between oxycodone-exposed or control litters at either timepoint (*p*’s > 0.050). **B.** (Left) Schematic depicting spatial learning paradigm. The “cloud-like” images show placement of the four odors. (Right) Mean ± SEM latency (seconds, s) to return to littermate goal box during training (trials 1-4) and at test. There was no difference in performance between oxycodone-exposed and control litters (p = 0.767). For both groups, there was a main effect of trial (p *=* 0.018) with pups in the fixed configuration performed significantly better at test than the alternating configuration (p *=* 0.036). **C.** Maternal separation-induced ultrasonic vocalizations (USVs) on PN8. (Left graph) Mean ± SEM peak USV frequency (kHz) during maternal separation before (Pre-) and after (Post-) a short reunion (grey column). Oxycodone-exposed pups emitted higher-frequency USVs overall relative to control pups (*p =* 0.002). (Center graph). Mean ± SEM number of USVs emitted during separation periods. Number of calls decreased over time (*p* < 0.001) for both groups with no difference observed between groups (*p =* 0.495). (Right graph) Mean ± SEM peak USV power (dB) emitted during separation periods. A significant interaction of Minute by Sex (p = 0.029) with a trending increase in power during the fourth and fifth minutes in the female oxycodone group failed to reach significance at individual minutes (all *p*’s > 0.050) and thus male and female data were combined. For the combined male and female data, a significant interaction between Treatment and Minute (all *p*’s < 0.050) revealed altered peak power over time for oxycodone pups although there were no significant pairwise difference (all *p*’s > 0.376).

## Statistics

All analyses were performed with scores averaged within litters (except where noted) using Prism GraphPad 8.0 analytical software (Graphpad, San Diego, CA v8.2-v8.4 for the MacOS and v8.3 for the Windows OS). Independent sample *t*-tests were used to assess differences in litter size and maternal behavior. Mixed factor ANOVAs were used to assess differences in pup weight, dam weight dam, and liquid intake. Sex was included in mixed factor ANOVAs for the neurodevelopmental tests, nociceptive thermal test, USV tests, and precipitated withdrawal behaviors. Due to small sample sizes, sex was not included as a factor when analyzing spatial navigation. Tests for outliers were conducted using the ROUT outlier test (Q=0.10%) and identified outliers only for the blood analyses.

## Results

### Dam Liquid Intake

Oxycodone did not alter dam liquid intake during the perinatal period (Figure 1B). A mixed factor ANOVA showed a significant increase (*p* < 0.001) in liquid intake primarily driven by increased drinking overall during lactation relative to the pre-breeding period. There was no effect of drug treatment (*p* = 0.707) but there was a significant Drug treatment by Day interaction (*p* = 0.042). Posthoc tests did not identify treatment differences on any day (all *p’s* > 0.237). Thus, Experimental dams self-titrated oxycodone intake throughout pregnancy and nursing, drinking the same as Controls.

### Dam Weight

Experimental dams showed similar weight gain as water-only Controls (Figure 1C). A mixed factor ANOVA (Drug treatment x Day) showed a significant increase (*p* < 0.001) in dam weight in both groups between the first (G1) and last (G21) day of gestation that was maintained throughout lactation (PN0-PN14; *p* < 0.001). There was no main effect of Drug treatment (*p =* 0.575) but there was a significant Drug treatment by Day interaction (*p* = 0.003). Posthoc tests did not find significant differences between oxycodone and controls on any day (all *p’s* > 0.050) with a trend at G0 (*p* = 0.089). Thus, maternal continuous oral oxycodone self-administration did not alter overall weight gain over pregnancy.

### Maternal Behavior

Maternal behavior was not altered by oxycodone intake (Figures 1D and 1E). Independent samples *t*-tests revealed no significant difference between Control and Oxycodone-exposed litters in the total duration or frequency of any observed behavior at either day (all *p*’s >.091; Table 3). Hence, in this model, maternal oral oxycodone self-administration did not affect maternal behavior.

**Table 3:** Results of Independent samples *t*-test comparing maternal care, nesting behavior, dam activity, and rough handling in Oxycodone-exposed and Control dams

|  | *t* ratio | df | *p* value |
| --- | --- | --- | --- |
| *Maternal Care* |  |  |  |
| Nursing on pups | 1.50 | 9 | 0.169 |
| Grooming pups | 0.44 | 9 | 0.671 |
| Anogenital lick pups | 0.60 | 9 | 0.566 |
| *Nesting Behavior* |  |  |  |
| Pups all in nest | 1.38 | 9 | 0.200 |
| Mom in nest | 1.89 | 9 | 0.091 |
| Nest building | 0.74 | 9 | 0.477 |
| *Dam Activity* |  |  |  |
| Mom active | 0.46 | 9 | 0.655 |
| Walking | 0.56 | 9 | 0.587 |
| Eat/drink | 0.73 | 9 | 0.483 |
| Rearing | 0.74 | 9 | 0.480 |
| Self-grooming | 0.45 | 9 | 0.660 |
| *Rough Handling* |  |  |  |
| Rough transport | 0.76 | 9 | 0.466 |
| Stepping on pups | 0.70 | 9 | 0.500 |
| Dragging pups | 0.16 | 9 | 0.878 |
| *Milk Ejections* | 1.15 | 9 | 0.279 |

**Note:** Maternal oxycodone consumption during and after gestation did not change maternal behavior relative to water-only control dams.

### Pup and Litter Measures

#### Litter size

Maternal oxycodone consumption did not alter Litter Size relative to water-only Controls (Figure 1F). An independent samples *t*-test revealed no significant difference in Litter Size (*p* > 0.999) between Control and Oxycodone-exposed dams prior to culling. Thus, maternal continuous oral oxycodone self-administration did not affect fetal viability.

#### Pup Weight

Prenatal oxycodone exposure had no effect on pup weight at birth or weight gain relative to Control pups during the postpartum period (Figure 1G). Mixed-factor ANOVA (Sex x Drug Treatment x Pup Age) revealed no main effect of Sex (*p* = 0.198) nor any significant Sex interactions (all *p’s* > 0.291). Pups gained weight over time (*p* < 0.001) between PN2 and PN14 with no significant effect of Drug Treatment (*p* = 0.605) or Age by Treatment interaction (*p* = 0.978). Therefore, maternal continuous oral oxycodone self-administration did not affect the offspring growth rate.

### Blood Levels

Data from males and females were combined. Oxycodone was detectable in blood at all ages (Figure 1H). There were two aberrant blood assay results that were identified by the ROUT outlier test (Q=0.10%). These were one dam (130 ng/mL) and one PN14 pup (57 ng/mL), and those two data points were excluded. A one-way ANOVA (Age including Dams) found no significant change in blood levels over age (*p* = 0.069), with the caveat of a limited sample size. These data suggest that oxycodone is passed from the lactating mom to the pups, albeit pups’ levels are lower than maternal levels.

### Neurodevelopmental Tests

Neurobehavioral outcomes were not altered by maternal oxycodone (all *p*’s > 0.050). Therefore, there were no gross impairments in sensory-motor function due to drug exposure throughout early development (Figure 2, Table 4).

**Table 4.** Mixed-Factor ANOVA analyses of neurodevelopmental and thermal response test between oxycodone-exposed and control pups

| PN3 |  |  |  |  | PN7 |  |  |  |
| --- | --- | --- | --- | --- | --- | --- | --- | --- |
|  |  | DFn,DFd | F ratio | p value |  | DFn,DFd | F ratio | p value |
| *Geotaxis* | Drug x Sex | 1, 15 | 0.04 | 0.853 |  | 1, 15 | 0.04 | 0.853 |
|  | Drug | 1, 15 | 0.14 | 0.716 |  | 1, 15 | 0.14 | 0.716 |
|  | Sex | 1, 15 | 0.01 | 0.920 |  | 1, 15 | 0.01 | 0.920 |
| *Righting* | Drug x Sex | 1, 15 | 0.71 | 0.414 |  | 1, 14 | 0.24 | 0.630 |
|  | Drug | 1, 15 | 0.19 | 0.673 |  | 1, 14 | 0.48 | 0.498 |
|  | Sex | 1, 15 | 0.27 | 0.608 |  | 1, 14 | 0.01 | 0.923 |
| *Cliff Avoidance* | Drug x Sex | 1, 15 | 0.05 | 0.823 |  | 1, 14 | 2.39 | 0.144 |
|  | Drug | 1, 15 | 2.71 | 0.121 |  | 1, 14 | 0.00 | 0.971 |
|  | Sex | 1, 15 | 0.15 | 0.702 |  | 1, 14 | 0.04 | 0.847 |
| *Pivoting* |  |  |  |  |  | 1, 14 | 0.01 | 0.916 |
|  |  |  |  |  |  | 1, 12 | 3.95 | 0.070 |
|  |  |  |  |  |  | 1, 12 | 0.04 | 0.850 |
| PN2 |  |  |  |  | PN14 |  |  |  |
| *Thermal Test* | Drug x Sex | 1, 12 | 0.41 | 0.533 |  | 1, 25 | 0.04 | 0.841 |
|  | Drug | 1, 12 | 1.58 | 0.233 |  | 1, 25 | 2.08 | 0.162 |
|  | Sex | 1, 12 | 0.35 | 0.563 |  | 1, 25 | 0.00 | 1.000 |

**Note:** Oxycodone exposure did not alter pup development or response to thermal stimuli.

### Nociceptive Thermal Tests

Experimental and Control pups were similar in response to a thermal stimulus at the early (PN2) and later (PN14) stages of development (all *p’s* > 0.160; Figure 3A, Table 4). This suggests that oxycodone blood levels in pups were not sufficient to elicit tolerance or hyperalgesia even though oxycodone was continuously available to the mother.

### Spatial Navigation

When separated by Sex, N’s were small (Oxycodone n=2-3, Control n=4-5) so all analyses are collapsed across sex. Both Control and Oxycodone-exposed litters showed spatial learning in the Fixed, but not Alternating, configuration (Figure 3B). Perinatal oxycodone exposure did not affect hippocampus-dependent spatial learning (Figure 3B). There was no effect of Drug (*p* = 0.767) but there was an effect of Trial (*p* < 0.001) and Configuration (Alternating vs. Fixed, *p* = 0.018). The latency to find the goal box decreased from the first training trial to the test trial for both Control and Oxycodone-exposed litters (*p*’s = 0.036) with shorter latencies observed for animals trained using the fixed configuration relative to the alternating configuration. Thus, performance improved as a function of learning over training trials in the fixed configuration compared to the alternating configuration. There were no interactions of Trial by Drug treatment (*p* = 0.939), Trial by Configuration (*p* = 0.154), Drug Treatment by Configuration (*p* = 0.950) or three-way interaction (*p* = 0.829).

### Maternal Separation-induced Vocalizations

Perinatal oxycodone exposure increased USV peak frequency and altered power over time but did not affect call number at PN8 both before and after a two-minute separation from the mother (Figure 3). There was no effect of pre-versus post-separation (all *p*’s > 0.050) so all results are collapsed across separation.

#### Peak Frequency

There was no effect of sex on USV peak frequency (all *p’s* > 0.103) so results are collapsed across peak frequency. Vocalization peak frequency increased in all pups over the course of the five minute separations (p < 0.002), and perinatal oxycodone exposure significantly increased vocalization peak frequency relative to Controls (p = 0.031) There was no interaction between Drug Treatment and Minute (p = 0.438).

#### Number of USV’s

There was no effect of sex on number of calls emitted (*p* = 0.342) so results are collapsed for number of USVs. The number of calls emitted in both oxycodone and control animals decreased across time (*p* < 0.001); however, there was no difference in the number of calls between groups (*p* = 0.496) and no interaction between Time and Treatment (*p* = 0.438).

#### Peak Power

There were significant Minute by Sex (*p* = 0.029) and Minute by Treatment by Sex (*p* = 0.031) interactions. *Post hoc* analysis revealed no significant pairwise differences (all *p*’s > 0.050). Subsequent analysis of males and females separately revealed significant Minute by Treatment interactions, (*p* = 0.036) and (*p* = 0.023), respectively. Post hoc analysis did not show significant pairwise Minute differences between Oxycodone- and Control-groups (all *p*’s > 0.376). Hence, maternal continuous oral oxycodone self-administration induced changes in affective behaviors as judged by an increased peak frequency and differences in power over time of pup calls.

### Precipitated withdrawal

Overall, oxycodone-exposed animals increased activity relative to Controls regardless of antagonist treatment (naltrexone or saline, Figure 4A). There were no effects of sex (all *p*’s > 0.050), so all analyses are collapsed across sex (Table 5).

**Table 5.** Mixed factor ANOVA analyses of pup behavior in oxycodone-exposed and control pups following i.p. injections of naltrexone on PN9.

|  |  | Total Duration (s) |  |  | Total Frequency |  |  |
| --- | --- | --- | --- | --- | --- | --- | --- |
|  |  | DFn, DFd | F ratio | p value | DFn, DFd | F ratio | p value |
| *Face Washing* | Perinatal Drug Exposure | 1, 27 | 5.29 | **0.029** | 1, 27 | 3.25 | 0.082 |
|  | Naltrexone/Saline | 1, 27 | 0.03 | 0.857 | 1, 27 | 1.47 | 0.237 |
|  | Perinatal Drug Exposure x Naltrexone/Saline | 1, 27 | 1.09 | 0.307 | 1, 27 | 0.00 | 0.982 |
| *Walking* | Perinatal Drug Exposure | 1, 27 | 3.42 | 0.076 | 1, 14 | 4.10 | 0.063 |
|  | Naltrexone/Saline | 1, 27 | 0.00 | 0.979 | 1, 13 | 0.70 | 0.418 |
|  | Perinatal Drug Exposure x Naltrexone/Saline | 1, 27 | 1.00 | 0.325 | 1, 13 | 0.21 | 0.652 |
| *Moving Head and Paws* | Perinatal Drug Exposure | 1, 27 | 8.15 | **0.008** | 1, 27 | 5.67 | **0.025** |
|  | Naltrexone/Saline | 1, 27 | 0.53 | 0.471 | 1, 27 | 1.43 | 0.242 |
|  | Perinatal Drug Exposure x Naltrexone/Saline | 1, 27 | 0.46 | 0.503 | 1, 27 | 0.00 | 0.960 |
| *Head Moves* | Perinatal Drug Exposure | 1, 27 | 0.29 | 0.596 | 1, 27 | 0.00 | 0.978 |
|  | Naltrexone/Saline | 1, 27 | 0.06 | 0.806 | 1, 27 | 0.58 | 0.454 |
|  | Perinatal Drug Exposure x Naltrexone/Saline | 1, 27 | 1.79 | 0.192 | 1, 27 | 0.68 | 0.415 |
| *Quiet* | Perinatal Drug Exposure | 1, 14 | 5.95 | **0.029** | 1, 14 | 2.941 | 0.108 |
|  | Naltrexone/Saline | 1, 13 | 0.01 | 0.916 | 1, 13 | 0.3416 | 0.569 |
|  | Perinatal Drug Exposure x Naltrexone/Saline | 1, 13 | 0.62 | 0.446 | 1, 13 | 0.2786 | 0.607 |

**Note:** Analyses are presented for behavior total duration (seconds, s) and total frequency. Oxycodone exposure increased active behaviors and decreased quiet regardless of naltrexone injection. Significant effects are represented with bolded text. pw drug=precipitated withdrawal drug.

**Figure 4:**
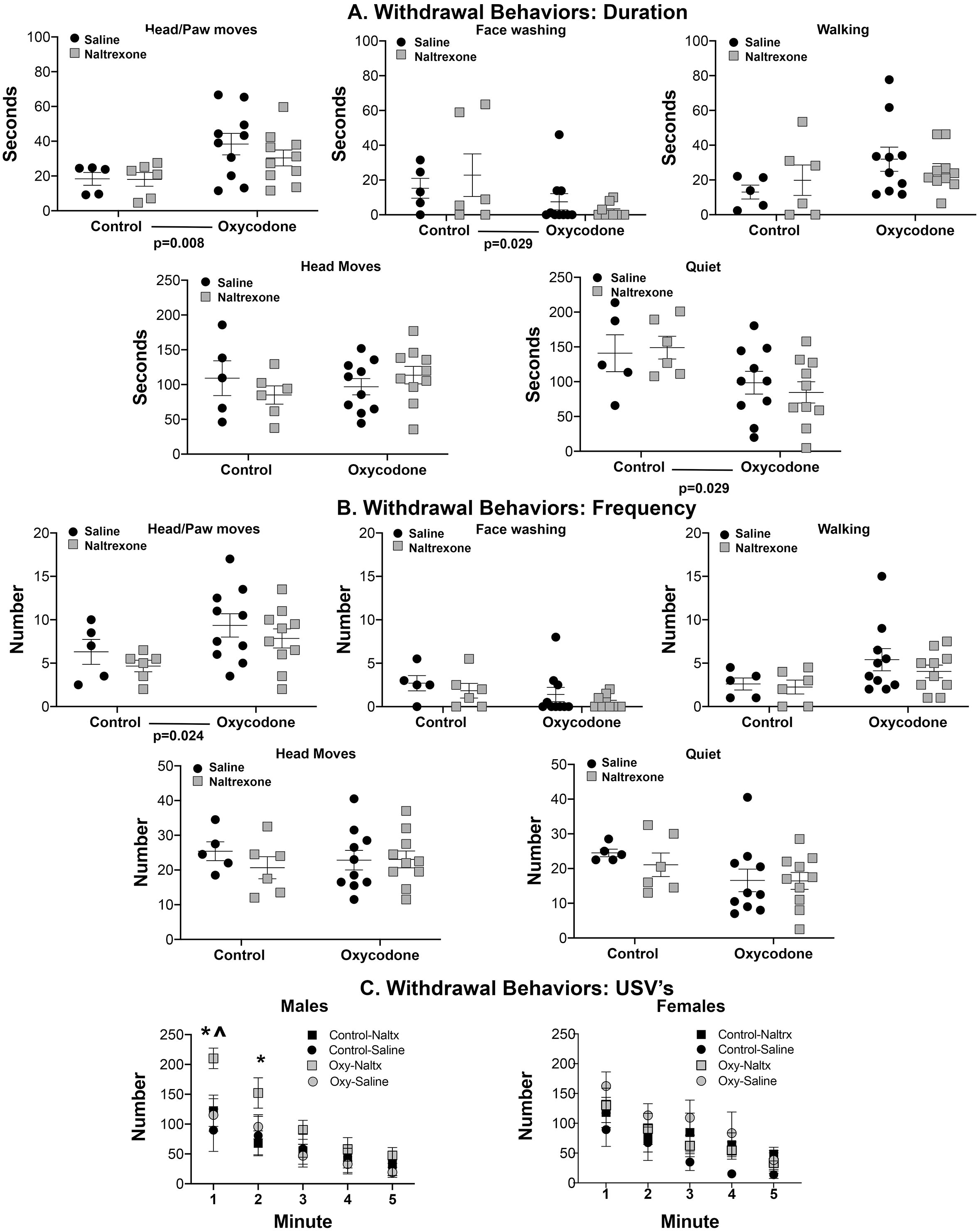
Mean ± SEM of naltrexone-precipitated withdrawal behaviors, (**A**) duration and (**B**) frequency, and USVs **(C)** on PN9. There were no effects of naltrexone treatment for any behavior (all *p*’s > 0.050). **A-B.** Oxycodone exposure increased concurrent head and paw movement total duration (*p =* 0.008) and frequency (*p =* 0.025). Face washing total duration (*p =* 0.029) was decreased in oxycodone-exposed pups relative to controls; however, frequency did not differ significantly between groups (*p =* 0.082). Walking in oxycodone pups showed non-significant trends toward increased total duration (*p =* 0.076) and frequency (*p =* 0.063). Head movement did not differ between groups in total duration (*p =* 0.806) or frequency (*p* = 0.454). Total duration of sitting quietly was increased (*p =* 0.029) in control pups relative to oxycodone-exposed pups. **C.** Mean ± SEM number of USVs. (Left graph) In males, for both groups cries decreased between the first and fifth minute (*p* < 0.001). However, naltrexone-injected oxycodone male pups emitted more cries relative to all other groups during the first minute (*p*’s < 0.050). (Right graph. For females, cries decreased between the first and fifth minute (*p* < 0.001) with no other effects observed (all *p*’s > 0.124). * = *p* < 0.050, Saline-naltrexone vs. Oxycodone-naltrexone; ^ = *p* < 0.050, Oxycodone-naltrexone vs. Oxycodone-saline.

#### Head and paw movements

Oxycodone exposure increased the total duration (*p* < 0.008) and frequency (*p =* 0.024) of concurrent head and paw movements regardless of naltrexone versus saline treatment.

#### Face washing

Face washing duration was decreased in Oxycodone-exposed pups relative to Controls (*p* = 0.029), although its frequency was not significant (*p* = 0.082).

#### Walking

There was no significant difference in total duration (*p* = 0.076) or frequency (*p =* 0.063) of pup walking.

#### Head Movements

There were no effects or interactions of perinatal treatment or drug on head movement total duration (*p*’s > 0.192) or frequency (all *p’*s >0.415).

#### Quiet time

Control pups spent significantly more time quiet relative to oxycodone-exposed pups (*p =* 0.029) regardless of drug treatment, although the number of quiet bouts did not differ (*p* = 0.108).

These data suggest that this model of maternal continuous oral oxycodone self-administration induces measurable levels of withdrawal behaviors in the offspring that are not further precipitated by opioid antagonist administration.

#### Ultrasonic vocalizations

##### Number of Vocalizations

Following naltrexone injection, the number of USVs increased in male but not female pups in the oxycodone group, although peak frequency and peak power were unchanged (Figure 4). In males, a three-way ANOVA for number of cries revealed a significant main effect of Minute (*p* < 0.001), an interaction of Minute by Drug treatment (*p* = 0.026) and Minute by Drug treatment (*p* = 0.012). Although the number of cries decreased generally between the first and fifth minute (*p* < 0.001), naltrexone-injected oxycodone pups emitted more cries relative to all other groups during the first minute (*p*’s < 0.050) and relative to Control-Naltrexone pups during the second minute (*p* = 0.020), with no differences observed between other groups at any time (*p*’s > 0.25). There were no other main effects or interactions (*p*’s > 0.050). In females (Figure 4), there was a significant main effect of Minute (*p* < 0.001) such that cries decreased between the first and fifth minute (*p* < 0.001); however there were no other main effects or interactions observed (all *p*’s > 0.124).

##### Peak Frequency and Power

Peak power and peak frequency analyses showed no main or interaction effects of naltrexone injection on USV peak power or frequency in males or females regardless of perinatal drug exposure (all *p*’s > 0.050 for power and all *p*’s > 0.115 for frequency). Therefore, only male offspring showed precipitated withdrawal effects following maternal oxycodone exposure.

## Discussion

Pre- and postnatal exposure to the semi-synthetic opioid oxycodone disrupted pup affective behavior while leaving neurodevelopment capabilities and spatial learning largely intact. This was not due to stunted growth or impoverished maternal care, as these were unaffected by perinatal oxycodone exposure.

### Oxycodone does not affect pup growth and has minimal effects on dam weight gain

Pup weight, litter size, and dam weight between control and oxycodone litters did not differ throughout gestation and nursing, with the exception of the day of parturition when oxycodone dams weighed more than controls. The current data largely corroborate previous oxycodone studies: prenatal oxycodone produces little to no effects on fecundity, gestation, or neonatal growth, although some variability exists across exposure protocols^22-25^. Similarly, maternal intra-atrial/venous oxycodone did not change litter size, sex distribution, or pup weight. It did, however, reduce maternal weight gain, which may be attributed to intermittent drug access and the confounding effects of withdrawal between drug exposures^5,24,25^. Oxycodone delivered by oral gavage, initiated prior to breeding, resulted in reduced birth weight in oxycodone-exposed pups that resolved by the first postnatal week^22^; however, if gavage was initiated during gestation, weight reductions persisted into adolescence^23^. Increased mortality and long-term reductions in pup weight after birth have been observed using other opioids, such as morphine, buprenorphine and methadone^6,7,10,11,13^; the reason for variations in opioid effects across drug and exposure protocols remains unclear. Likely when drug treatment is initiated prior to gestation and maternal withdrawal and stress are minimized, somatic effects on pups are minimized^16-18^. Additional experiments closely evaluating withdrawal scores in different maternal oxycodone exposure protocols are necessary to test this hypothesis.

### Maternal continuous oral self-administration oxycodone preserves typical maternal behavior

Maternal behavior was unaffected by pre-and perinatal ingestion of oxycodone. Acutely, endogenous opioids regulate the onset, modulation, and maintenance of maternal behavior^8,39-41^. The effects of chronic exposure are less clear. Oxycodone IV self-administration prior to and during gestation increased pup retrieval latency^5^. However, because no other measures of maternal behavior were analyzed, the full effect of this manipulation on maternal behavior is not known. Few reports measure maternal behavior for opioids other than morphine^14,22,24,25,42^ making the available information of their effect on maternal behavior limited. Morphine injections during gestation decreased pup-directed behaviors such as pup grooming, while increasing non-maternal activities such as dam self-care^8,40^, although effects on pup retrieval are not consistently observed^39,40^. These behavioral changes described are confounded with increased maternal stress due to drug initiation in the middle of or after gestation^41^. Also, behavioral changes due to opioid withdrawal cycles in studies of intermittent drug exposure muddle putative opioid effects. Indeed, differences in grooming and licking behavior are reduced if dams exposed to morphine during gestation are not exposed to morphine during lactation^39,40^. Moving forward, it will be important to control for withdrawal effects in such studies.

Clinical presentation and preclinical models of NOWS show impaired somatosensory development, altered affective behavior, and physiological abnormalities^6,9-11,13,25^. Rodent models using intermittent access, expose pups to drug peaks and *in utero* withdrawal^5-10,22-24,42^, causing poor fetal outcomes independent of specific drug effects^16^. Chronic exposure models (minipumps, pellets) prevent *in utero* withdrawal, but if initiated during gestation can cause maternal stress^17-19^. Moreover, pups experience withdrawal at birth if the maternal opioid is discontinued or pups are cross-fostered to non-exposed surrogates^12-15^. Here, using a continuous (24/7) oral self-administration paradigm, dams had access to oxycodone prior to conception and through lactation and could titrate their intake, reducing the likelihood of spontaneous withdrawal^27^, thus dissociating the consequences or drug exposure from withdrawal for the offspring. These parameters more closely mimic human fetal exposure both when the mother is using opioids illicitly and in clinical practice where mothers are maintained on opioid assisted programs to avoid the adverse effects of withdrawal on the newborn^1-4^.

### Maternal continuous oral self-administration of oxycodone does not impair infant spatial learning

We interrogated cognitive function in 13-day-old pups and found that both control and oxycodone-exposed animals performed the spatial learning task at PN13 similarly. The task is sensitive to perinatal insults early in development, at least in mice; perinatal exposure to environmental pollutants impaired olfactory-spatial learning in 11-day-old mice^38^. Others report that impairments can appear later in development with prenatal opioid exposure impairing spatial learning from adolescence^43^ to adulthood^22^. During spatial navigation tasks, adult mice prenatally-exposed to oxycodone took longer to find and traveled a greater distance to locate the hidden platform when long (40-50 minute) inter-trial intervals were used^22^. This oxycodone-related deficit was not observed using short intervals between trials (15-30 minute), perhaps suggesting a memory retention deficit over a long interval, implicating hippocampal dysfunction^44^. This lack of an oxycodone-related deficit is consistent with our present findings. In addition, when opioid exposure was restricted to the gestation period the pups underwent withdrawal thereafter^22,43^. In our study, pups were exposed to oxycodone through the mom at the time of testing, and this likely lack of withdrawal in our study possibly contributed to the pups in the present study not demonstrating a spatial learning deficit.

### Maternal continuous oral self-administration oxycodone disrupts pup affective behavior

Perinatal oxycodone significantly increased pup USV peak frequency during periods of pup separation from their mothers on PN8. Pup isolation-induced USVs are considered a measure of distress, a proxy for maternal attachment dampened by maternal presence, and critically-dependent on the developing opioid system^36,45^. Similar to our results, oxycodone exposure via maternal intravenous self-administration did not change total number of calls^25^. However, we also report the novel finding that perinatal oxycodone resulted in increased peak frequency and dampened decline in power of USV’s during maternal separation relative to control pups. Previous reports that show isolation-^45^ and stress-induced-^46^ shifts in USV frequency are indicative of a more negative affective state. Indeed, non-opioid drugs (ethanol) also produce shifts in isolation-induced USVs toward higher frequencies^47^. Together, these results suggest increased negative affect in pups following perinatal oxycodone exposure.

### Withdrawal behavior is differentially-expressed in oxycodone-exposed pups

On PN9, both saline- and naltrexone-injected oxycodone-exposed pups increased withdrawal behaviors relative to controls; however, only naltrexone-oxycodone male pups increased USVs. Pup withdrawal behaviors are distinct from those in adults and change from infancy through adulthood in the rat^11,20,48^. Chronic oxycodone exposure resulted in increased infant withdrawal behaviors relative to water controls, but naltrexone- and saline-treated pups perinatally exposed to oxycodone did not differ in withdrawal behaviors. One possible explanation is that saline-oxycodone animals could not be further precipitated with naltrexone because pups were already experiencing spontaneous withdrawal (e.g. oxycodone intake vs. water intake).

In contrast, naltrexone increased USVs in male pups. USV numbers, an affective measure of withdrawal sensitive to naltrexone treatment, were elevated in males only for the first two minutes after precipitated withdrawal. This is consistent with prior data where withdrawal was precipitated in pups injected twice daily with morphine^48^ or born of mothers implanted with methadone pellets^11^. Interestingly, females did not increase USVs although there are sex - dependent effects in oxycodone pharmacokinetics^49^. Our previous study in adult oxycodone oral self-administration showed how blood levels greatly differed between sexes^27^ with males showing lower blood levels when similar drug concentrations were ingested.

### Conclusion

Overall, our preclinical study recapitulates key aspects of chronic oral oxycodone self-administration in pregnant women and consequent neonatal withdrawal symptoms. Exposure to oxycodone as conducted in this study caused measurable detrimental effects on affective domains in developing pups. These effects are strikingly comparable to those observed clinically (i.e. high-pitched crying and irritability), suggesting that this route and mode of administration may be complementary to the already-existing opioid exposure models and can be utilized to investigate the immediate and long-term effects of oral self-administration on affect and cognition. Oral self-administration of oxycodone had fewer developmental effects than did studies giving other opioids through different routes and time windows of exposure, suggesting this route of oxycodone delivery targets developing affective systems more strongly than somatic, cognitive, and motor systems. The importance of our findings are two-fold: 1) We recapitulated aspects of neonatal opioid withdrawal syndrome and showed that maternal oral self-administration alters pup affective behavior, similar to what is observed in humans; and 2) Maternal oral self-administration of oxycodone may provide a suitable approach to reliably model the clinical scenario and further study the long-term trajectory of these altered affective components.

## Acknowledgements

We thank Ganesh S. Moorthy, Christina M. Vedar, and Athena F. Zuppa of CHOP’s Bioanalytical Core Center for Clinical Pharmacology for oxycodone analysis in blood, Peter Lenchur for help analyzing the USV data.

## Data Availability

The data that support the findings of this study are available from the corresponding authors upon reasonable request.

## Authors contributions (using CREDiT and COPE guidelines)

Conceptualization: GAB, AJE, GZ.

Methodology: GAB, AJE, MJD, PAR-D, GZ.

Software: Not applicable.

Validation: GAB, AAD, PAR-D, MJD, GZ.

Formal Analysis: GAB, AAD, PAR-D, GZ.

Investigation: GAB, AAD, HMD, AJE, PAR-D, DT, AV, GZ.

Resources: GAB, AJE.

Data Curation: GAB, AAD, MJD, DT, AV, GZ.

Writing - Original Draft: GAB, PAR-D, GZ.

Writing - Review & Editing: GAB, PAR-D, AAD, DT, AJE, RMS, AV, GZ.

Visualization: GAB, PAR-D, AAD, GZ.

Supervision: GAB, AJE, RMS, GZ.

Project Administration: GAB, AJE, GZ.

Funding Acquisition: GAB, AJE, RMS.

